# Seeing touch enhances the perception and processing of digitized gentle stroking

**DOI:** 10.1101/2025.11.13.688063

**Authors:** Brais Gonzalez Sousa, Daniel Senkowski, Shu-Chen Li

## Abstract

Observing touch activates brain regions similar to those activated by experiencing actual touch, suggesting that visual information can cross-modally influence tactile perception. In this electroencephalography (EEG) study, we investigated how viewing visual displays of an arm being touched may affect the perception and processing of digitally rendered touch patterns designed to resemble either stroking or tapping. Thirty-one participants experienced touch patterns delivered to their left forearm via a wearable sleeve while viewing either a photo of an arm or spatiotemporally aligned videos of an arm being touched in synchrony with either of the two touch patterns. Continuity and pleasantness ratings of touch stimuli were higher for stroking than for tapping. Correlations between continuity and pleasantness ratings were stronger when touch was accompanied by videos of touch than by the photo of an arm. Analysis of evoked brain responses revealed visual modulation of touch processing at centroparietal electrodes beginning at around 0.9 s, with the cross-modal effects diverging between stroking and tapping at about 1.6 s. Furthermore, the interaction effects of cross-modal influences between stroking and tapping at the neural level positively correlated with the visual modulation of pleasantness ratings in two right frontal clusters at around 1.4 s and 1.8 s. These results suggest that observing touch influences the perception and processing of touch through initial sensory integration at centroparietal sites, followed by later frontal valuation processes. This extends previous findings on affective touch by demonstrating that visual inputs can cross-modally shape the hedonic evaluation of digitally actuated touch.

## 1 Introduction

Perceiving a touch by another person is inherently multisensory and can convey different social messages that the receiver can categorize and interpret. Of particular interest here are two kinds of touch. Slow, gentle stroking is typically perceived as pleasant and reflects social-affective intentions. Tapping, however, is mostly perceived as attention-seeking and emotionally more neutral (Etzi et al., 2018; Löken et al., 2009; McIntyre et al., 2022; see Morrison et al., 2010 for a review). While stroking and tapping recruit distinct neural processes, contextual factors, like inputs from other sensory modalities, may also modulate neural responses and thereby shape how touch is perceived and interpreted (see Ellingsen et al., 2016 for review).

Electrophysiological and functional neuroimaging studies have revealed that both pleasant and neutral touch stimuli initially activate the primary somatosensory cortex (S1) but later diverge in terms of their temporal dynamics and the neural networks they engage. The S1 is engaged in processing both types of touch (Morrison, 2016), with its activity observable within the first 100 ms reflecting the rapid detection of tactile stimuli (Auksztulewicz et al., 2012; Eriksson Hagberg et al., 2019). Slow stroking also elicits a later negative amplitude peaking around 400 ms (sN400) that correlates with pleasantness ratings, indicating its role in affective processing (Schirmer, Lai, et al., 2022; Schirmer & McGlone, 2019). Beyond basic tactile detection, pleasant touch also activates the insular cortex (Björnsdotter et al., 2009; Morrison, 2016; Morrison et al., 2010; Olausson et al., 2002; Voos et al., 2013) associated with interoceptive perception (Craig, 2009) and the posterior temporal sulcus (Bennett et al., 2014; Bjornsdotter et al., 2014; Miguel et al., 2020; Voos et al., 2013) supporting social interactions (Frith & Frith, 2003; Isik et al., 2017; Yang et al., 2015). These signals relay to frontal regions, such as the orbitofrontal (Rolls et al., 2003; Voos et al., 2013) or medial prefrontal cortex (Gordon et al., 2013; Voos et al., 2013), for higher-order valuation processing. Electrophysiological studies further identify a late positive potential over the frontal cortex approximately 700 ms after touch onset (Ackerley et al., 2013; Schirmer, Lai, et al., 2022). This positive potential was localized to the superior frontal gyrus and frontal motor areas (Eriksson Hagberg et al., 2019), distinguishing affective from non-affective touch processing. Overall, pleasant touch involves a cascade of neural processing. This begins with early sensory encoding in S1, followed by activities in other regions processing socio-emotional features of the touch stimulus and valuating the experience.

Information from other sensory modalities that are not directly related to the touch stimuli, like odours (Croy et al., 2016) or pictures of facial expressions (Ellingsen et al., 2014), have been shown to also affect the perceived pleasantness of touch (Saarinen et al., 2021 for a review). However, it is still not well understood how informative cross-modal stimuli, like observing being touched, may influence the perception and processing of pleasant touch. Thus far, cross-modal visual influences on tactile perception have mainly been studied for noxious and neutral stimuli (Cardini et al., 2012; Eads et al., 2015; Höfle et al., 2012, 2013; Pomper et al., 2013). For noxious stimuli, visual information has been found to modulate local pain representations in sensorimotor areas and salience networks (Senkowski et al., 2014). Furthermore, participants reported greater unpleasantness when seeing a pain producing versus a neutral object approaching their body (Höfle et al., 2012). For neutral tactile stimuli, sensory acuity improved when participants were presented with photos of the stimulated body part (Eads et al., 2015), but not when they were presented with neutral objects (Cardini et al., 2012). This effect of ‘visual enhancement of touch’ (Kennett et al., 2001) was associated with larger N140 amplitudes at central electrodes, indicating increased processing of tactile stimuli in the somatosensory cortex (Taylor-Clarke et al., 2002). Cross-modal enhancement was also shown to involve the posterior parietal cortex supporting integration of visual and tactile signals (Nakashita et al., 2008; Pasalar et al., 2010).

While the neural mechanisms underlying cross-modal effects on noxious and neutral tactile stimuli have attracted more research attention, evidence on how visual stimuli affect pleasant touch has mainly focused on vicarious touch (i.e., seeing touch) without actual tactile stimulation. Perceived pleasantness of observed touch derived from unisensory visual information of touch exhibits similar velocity sensitivity to that of unisensory tactile touch; for example, observing slow stroking is perceived as more pleasant than observing fast stroking (Morrison, Bjornsdotter, et al., 2011). Furthermore, pictures of social touch interactions were perceived as inducing stronger positive emotions than inanimate or non-social control pictures (Peled-Avron et al., 2016). Despite not having actual tactile input, visual presentations of social touch interactions were found to activate the S1 (McCabe et al., 2008; Peled-Avron et al., 2016; Schirmer & McGlone, 2019). Interestingly, visual presentations of social touch interactions also showed a late positive potential when compared to pictures not involving touch (Schirmer & McGlone, 2019). The social components of the visually transmitted interactions are likely processed in association areas such as the posterior superior temporal sulcus or the temporo-parietal junction (Masson & Isik, 2023). These areas are also involved in visual motion processing (Kaiser et al., 2012) and understanding others’ mental states (Frith & Frith, 2003; Isik et al., 2017; Yang et al., 2015). Taken together, vicarious pleasant touch elicits behavioural and neural signatures that are comparable to those of unisensory pleasant tactile stimulation. However, the cross-modal modulation of sensory, socio-emotional or valuation processing of pleasant touch by informative visual stimuli remains to be investigated.

This electroencephalography (EEG) study builds on and extends previous research to examine the impact of visual information on the processing and perception of digitally rendered pleasant and neutral touch. Two types of touch, stroking and tapping, were delivered via a wearable shape-memory alloy (SMA) arm-sleeve (Muthukumarana et al., 2020). The SMA-based arm-sleeve enabled a precise synchronisation of digitally rendered tactile patterns with the visual stimulation. While human-to-human touch offers greater ecological validity (e.g., Gazzola et al., 2012), its temporal precision is limited and difficult to control. Conversely, robotic touch devices allow for more precise manipulation of stimulation parameters, but they are mechanically complex and may compromise ecological validity (e.g., Schirmer et al., 2022). The wearable arm-sleeve used here strikes a balance by providing better experimental control than human touch, while maintaining ecological validity by allowing a larger area of direct skin contact and aligning the physical arm position with the visual stimuli. Based on previous findings showing enhancing effects of cross-modal visual information (Eads et al., 2015; Höfle et al., 2012, 2013; Pomper et al., 2013), we predicted that adding informative visual inputs to the tactile stimulation would amplify differences in continuity and pleasantness ratings between conditions with digitally rendered stroking and tapping. We also expected that these behavioural effects would be accompanied by associated effects in cortical regions implicating multisensory perception of social touch, such as primary sensory, multisensory, and frontal areas.

## 2 Materials and methods

### 2.1 Participants

Forty-two young adults participated in this study and were reimbursed for their participation. Except for one participant, all were right-handed as assessed by the Edinburgh Handedness Inventory (Oldfield, 1971) and none reported clinical symptoms in the four weeks prior to the experiment. All participants provided written informed consent before the study, which was approved by the Ethics Committee of the TUD Dresden University of Technology (Approval number: SK-EK-5012021-Amendment). Eleven participants had to be excluded after EEG preprocessing and artefact correction/rejection because the number of trials after artefact rejection did not allow for calculation of reliable ERPs, i.e., their dataset contained less than 30 trials in at least one condition. These exclusions can be mostly attributed to sweating-related artifacts and head/body movement because of fatigue towards the end of the experiment. Thus, the effective sample consisted of 31 participants (mean (M) age = 23.58; standard deviation (SD) = 3.21 years; 18 women and 13 men).

### 2.2 Experimental setup

The experiment comprised two tactile conditions that manipulated touch types: stroking (T_Str_) and tapping (T_Tap_). The two touch types were accompanied by two visual conditions that manipulated the type of visual information: visual-video (V_V_) and visual-photo (V_P_). This resulted in a 2 x 2 within-subject design with four bisensory conditions: V_V_T_Str_, V_V_T_Tap_, V_P_T_Str_, V_P_T_Tap_. Additionally, two video-only (unisensory visual) conditions (V_V-Str_, V_V-Tap_) were presented to compare and identify brain activity unrelated to tactile stimulation (Fig. 1A). In the experiment, the participants’ task was to rate the continuity and pleasantness of the stimulation after bisensory trials (see Procedure).

**Fig. 1.**
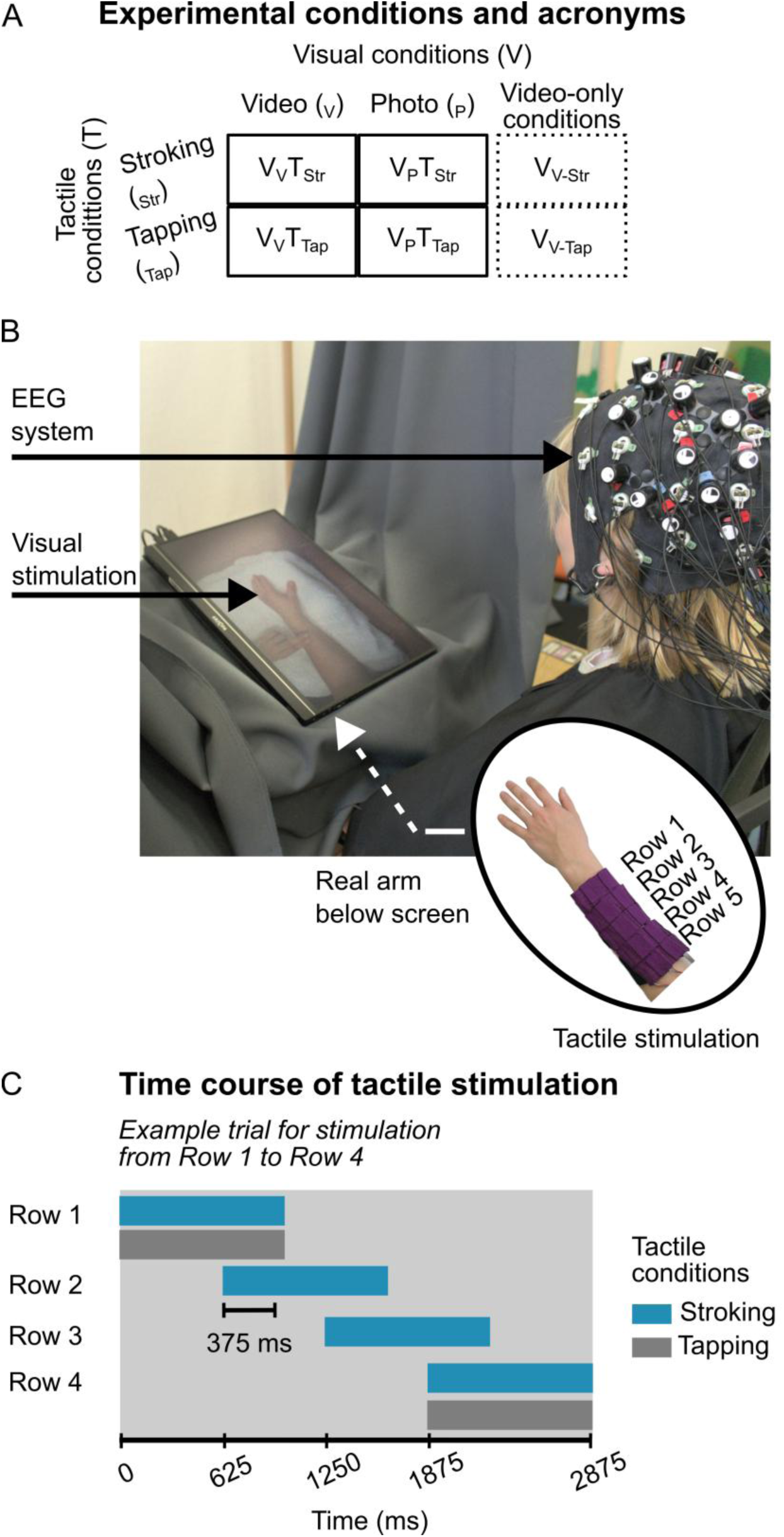
Experimental setup and conditions. (A) Tactile (stroking or tapping) and visual stimuli (video clips or a photo of a forearm) were combined to create four bisensory conditions (V_V_T_Str_, V_V_T_Tap_, V_P_T_Str_, V_P_T_Tap_). In addition, two video-only control conditions were presented (V_V-Str_, V_V-Tap_) (B) Each participant wore a wearable shape-memory alloy-based arm-sleeve on their left forearm that actuated two types of touch pattern: stroking or tapping. Participants placed their left arm below a flat screen, which displayed visual stimuli aligned with their left arm. They also wore a black cape to occlude them from seeing the wooden apparatus and their own forearm, while EEG data were recorded. (C) The time course of tactile actuations shows example trials for stroking (blue) and tapping (grey). Rows are numbered on the sleeve shown on the lower right of (A). Stroking was generated by consecutively actuating e.g., row 1 to 4 with a 375 ms overlap between rows, while for tapping only rows 1 and 4 were actuated with an 875 ms delay in between. Both stimuli lasted for 2.875 s.

### 2.3 Tactile stimuli

Slow stroking and tapping were digitally actuated and presented as tactile stimuli on the left forearm of the participant. These two touch patterns were generated with a shape-memory alloy-based arm-sleeve (see Fig. 1B for our setup and Srivastava et al., 2022 for an overview). Stroking was elicited by successively actuating across four rows on the sleeve resulting in a duration of 2.875 s at an average velocity of 4.17 cm/s. For tapping, the first and fourth row were actuated with a delay of 0.875 s in between, which also resulted in a trial duration of 2.875 s (see Fig. 1C). We previously showed that using a similar arm-sleeve and technical settings of stimulation intensity and duration, the stroking stimulation was rated more continuous yielded higher facial muscle activities associated with positive affect than the tapping stimulation (Sousa et al., 2024). We use the same duration for both touch types to avoid on-/offset related differences that may confound the analyses and results of EEG data. To reduce tactile habituation, the onset location and movement direction of the touch stimulations varied across trials (for details on the tactile device, see Supplementary material).

### 2.4 Visual stimuli

Visual stimuli were presented simultaneously with the touch onset elicited by the arm-sleeve. Two types of visual stimuli were used: videos of touch (stroking or tapping) or a photo of the forearm. The videos showed two fingers of a left hand performing strokes or taps on a left forearm resting at the same position and in the same orientation as the participants’ arm. The videos were recorded for the different variations of touch onset/movement direction, resulting in four clips per touch type. All clips lasted 2.875 s and were edited in Davinci Resolve (v. 17.4.6) for precise spatiotemporal alignment with tactile stimuli. Additionally, as a control of basic visual input of the touched body part, a photo of the left forearm not informative of being touched was displayed for the V_P_T_Str_ and V_P_T_Tap_ conditions (c.f. Kennett et al., 2001). We attempted to present fingers and an arm in the video clips and photos that were sex neutral.

### 2.5 Procedure

Each participant read instructions about the experiment and filled out the Edinburgh Handedness Inventory (Oldfield, 1971) before being seated in front of a wooden apparatus on which a 15.6 inch LED screen (141 pixels per inch, refresh rate: 144 Hz, contrast ratio: 1200:1, brightness: 300 cd/m²) was placed (Fig. 1B). Visual stimuli were presented at 120 Hz. For trials with tactile stimulation, the arm-sleeve was triggered via serial commands one second before the end of a fixation cross shown on the screen. This one-second interval corresponded to the latency needed to energize the shape-memory alloy (Muthukumarana et al., 2020).

Participants rested their left forearm comfortably below the screen on a Styrofoam pad wrapped in a towel with the arm-sleeve strapped onto their left dorsal forearm. For assessing brain correlates, an actiCAP was used to place EEG and near-infrared spectroscopy optodes. The current study focused explicitly on the analysis of behavioural and EEG data. During the experiment, pink noise was presented via earphones to mask the low-volume sound of the tactile device. The experiment ran on a custom-made script in PsychoPy (v2022.2.5, Peirce et al., 2019) that also sent serial commands to the arm-sleeve and triggered EEG acquisition. It entailed five blocks, each with 48 trials that were presented in a random order with eight trials per condition per block. The eight trials in each block of a given condition consisted of two repetitions of the four different variations of touch onset/movement direction. Overall, this resulted in 40 trials in each of the six conditions. On average the experiment lasted around 57 minutes (SD = 7.6 minutes).

Before the experimental blocks, each participant was presented with the position of the visually displayed virtual arm and was asked to align their left forearm with the arm displayed on the screen. Subjective embodiment was rated after completing all experimental blocks as a manipulation check. A trial started with a fixation cross with randomized durations of 2.5, 2.75, or 3 s. In bisensory trials, visual display and tactile stimulation started simultaneously and lasted for 2.875 s. After stimulus presentation, participants viewed a 0.5 s fixation cross, and only after bisensory trials, responded to two rating scales of pleasantness and continuity. In every block, at the end of three randomly selected trials, participant’s visual attention was assessed by the multiple-choice question, ‘Which video/photo was presented during the last trial?’ At the end of each block, participants were able to take a break with a self-determined duration.

### 2.6 Subjective ratings

For pleasantness ratings, participants responded to the short text prompt, ‘I felt the touch was…’, by using a continuous scale ranging from ‘unpleasant’ (−10) to ‘pleasant’ (+10). The continuity ratings were assessed via the statement, ‘The touch felt more like…’, by using a scale with endpoints labelled respectively as ‘tapping’ (−10) and ‘stroking’ (+10). Because both statements assess subjective perception of a ‘felt touch’, these ratings were only collected for bisensory trials and not for unisensory visual trials.

To validate that participants sufficiently felt they were looking at their own arm when they were viewing the video clips of touch and photos of an arm, we assessed the participant’s subjective feeling of embodiment of the visually displayed arm. For this, a modified 10-item questionnaire (Longo et al., 2008) was administered after completing all five experimental blocks. The questionnaire was translated to German and adapted for our study purpose (e.g., the term ‘rubber hand’ was substituted by ‘virtual arm’). It encompassed the subscales of ownership, location, and agency. The four variations (differing in start location and movement direction) of a condition were presented consecutively, with a brief 1-second fixation cross separating each presentation. Participants rated their experienced embodiment for each of the presented conditions on a continuous scale from ‘completely disagree’ (−10) to ‘completely agree’ (+10).

### 2.7 EEG data acquisition

EEG data were recorded with a 32-channel EEG system (LiveAmp; Brain Products) with active Ag/AgCl electrodes mounted according to the international 10-20 system and acquired via BrainVision Recorder (Version 1.25.0001; Brain Products). The FCz and FPz electrodes served as online reference and ground electrodes, respectively. EEG data were sampled at 500 Hz and online filtering consisted of a low-pass filter at 131 Hz with no high-pass filter (DC recording). The average impedance across 25 participants and all electrodes was 7.37 *k*Ω (SD: 7.09 *k*Ω). Due to a technical issue of the recording software the impedance measurements of 8 participants were not saved with the EEG data files but were kept below a predefined threshold of 25 *k*Ω prior to data collection.

### 2.8 EEG data preprocessing

EEG data were pre-processed offline using MNE-Python (Gramfort, 2013) in Python 3.11.1. First, two zero-phase finite-impulse filter were applied to the EEG data with a low-pass filter at 30 Hz (7.5 Hz transition bandwidth, −6dB cut-off) and a high-pass filter at 0.5 Hz (0.5 Hz transition bandwidth, −6dB cut-off). Noisy data channels were interpolated, and the interpolated EEG data were re-referenced to the channel average and epoched from −1 s to 2.875 s around the stimulus onset, and downsampled to 250 Hz via decimation. The data were then manually examined, and epochs containing unusual artifacts (e.g., drifts, muscle movements) were excluded.

For the subsequent independent component analysis (fastICA, 1000 iterations) for artefact rejection, a copy of the cleaned EEG data was created with the high-pass filter increased to 1 Hz while keeping the low-pass filter at 30 Hz. The copied EEG data were subjected to the fastICA and the resulting component structure from the 1-30 Hz data was applied to the preprocessed data which had been filtered at 0.5-30 Hz. Components reflecting artefacts, i.e., eye movement, eye blinking, channel or line noise, were manually inspected and removed. Subsequently, data were back-projected from the component space to EEG space without the removed artefacts. Epochs in the corrected EEG space underwent a final visual inspection and epochs containing residual artefacts were excluded. On average 216 (90%) epochs (SD: 14 epochs) remained for the EEG data analyses.

### 2.9 Statistical analysis

#### 2.9.1 Behavioural analysis of continuity, pleasantness and embodiment subscales

To investigate cross-modal visual effects on behavioural ratings of continuity and pleasantness, we analysed tactile and visual conditions (see Fig. 1A), as well as their interaction as fixed effects in two separate linear mixed models (LMMs). Because previous studies have reported sex/gender differences (e.g., Schirmer, Cham, et al., 2022) and effects of ageing (e.g., Sehlstedt et al., 2016) on affective touch perception, both LMMs included gender and age as covariates. The random effect structure was maximal with a random slope per participant for the interaction effect of tactile by visual conditions.

To investigate the question of touch type on subjective feeling of the embodiment, we analysed ratings of the embodiment subscales, i.e., location, ownership, and agency, assessed in bisensory conditions presented with videos. Comparability with previous research (e.g., Höfle et al., 2012) was ensured by rescaling ratings between 1 and 6. The average of the three subscale ratings were modelled in separate LMMs with tactile conditions as fixed effects, and an intercept per subject as random effects. Additionally, gender and age were included as fixed effects to investigate potential gender differences and age effects on the embodiment subscales. To evaluate the significance of fixed effects, F-tests were performed for all LMMs using Satterthwaite approximation for degrees of freedom at an alpha threshold of .05. Post-hoc t-tests tested differences between estimated marginal means (EMM) of the significant fixed effects while correcting for multiple testing via Tukey adjustments (Luke, 2017). Effect sizes (*d*) for LMMs were calculated based on the formula presented by Westfall et al. (2014).

#### 2.9.2 Analysis of event-related potentials

For the analysis of event-related potentials (ERPs), epochs were baseline corrected using the mean voltage in the time-window from −0.3 to 0 s before to the onset of tactile or visual stimuli. The first step of the ERP analysis was taken to establish whether the digitally delivered tactile stimuli resulted in prominent ERP components over the contralateral somatosensory cortex (Katus & Eimer, 2019), regardless of the cross-modal modulation. Specifically, tactile processing was verified by the presence of a central-contralateral component (N2cc) at C4, i.e., the contralateral somatosensory cortex. The initial stroking and tapping amplitudes in the visual-photo condition were averaged from 0.18 to 0.28 s and compared between C3 (ipsilateral somatosensory cortex) and C4 (Katus & Eimer, 2019). ERPs were modelled using a LMM with tactile condition (V_P_T_Str_ versus V_P_T_Tap_) and laterality (C3 versus C4) as fixed effects, and an intercept per subject as a random effect.

The next step of the ERP analysis focused on cross-modal visual effects (i.e., a video showing touch actions versus a photo of an arm) on stroking or tapping. These effects were analysed using spatiotemporal cluster-based permutation tests (Maris & Oostenveld, 2007). ERP differences between video and photo conditions were separately tested for stroking (V_V_T_Str_ - V_P_T_Str_) and tapping (V_V_T_Tap_ - V_P_T_Tap_) at each sampling point using two-tailed paired t-tests with a cluster-forming threshold of .05. The spatiotemporal clusters formed in these analyses were then tested against a null distribution obtained from Monte-Carlo sampling at 1000 permutations using a Bonferroni-corrected alpha threshold of α_Bonferroni_ = .05/2 = .025. Furthermore, we were interested in how visual input may affect tactile stimulation differently. To this end, we tested spatiotemporal clusters of ERPs calculated as the interaction between the visual and tactile conditions (i.e., (V_V_T_Str_ - V_P_T_Str_) – (V_V_T_Tap_ - V_P_T_Tap_)) in a spatiotemporal cluster-based permutation test (two-tailed paired t-test, cluster-forming threshold = .05) against a null distribution obtained from Monte Carlo sampling at 1000 permutations.

Finally, we investigated for each touch type whether differences in the cross-modal processing of videos versus photo in ERPs were associated with cross-modal processing differences in pleasantness ratings. This analysis was done to examine the relationship between cross-modal effects on neural processing and the subjective behavioural ratings of pleasantness. For each participant, pleasantness ratings were averaged across trials for each tactile condition (stroking, tapping) to calculate the difference between the videos and the photo condition. Spatiotemporal cluster-based permutation tests assessed correlations between ERP differences and pleasantness rating differences separately for stroking (ERPs: V_V_T_Str_ - V_P_T_Str_; pleasantness ratings: V_V_T_Str_ - V_P_T_Str_) and tapping (ERPs: V_V_T_Tap_ - V_P_T_Tap_; pleasantness ratings: V_V_T_Tap_ - V_P_T_Tap_) conditions (Spearman’s rank correlation, cluster-forming threshold = .01). A cluster indicated a significant correlation if the cluster-level statistic exceeded the corrected critical alpha of α_Bonferroni_ = .05/2 = .025 in a null distribution obtained from Monte-Carlo sampling at 1000 permutations.

We also investigated whether neural differences based on the interaction contrast between tactile and visual conditions were correlated with the interaction in pleasantness ratings between those conditions. This interaction contrast ((V_V_T_Str_ - V_P_T_Str_) – (V_V_T_Tap_ - V_P_T_Tap_)) was calculated for ERPs and pleasantness ratings. Significant correlations between ERP and pleasantness interactions were obtained from a spatiotemporal cluster-based permutation test (Spearman’s rank correlation, cluster-forming threshold = .01) where the cluster statistic was tested against a Monte-Carlo sampled null distribution at 1000 permutations.

## 3 Results

### 3.1 Behavioural ratings of continuity, pleasantness, and embodiment subscales

Continuity ratings showed a significant main effect for tactile conditions (*F*_1,30_ = 193.71, *p* < .001, *d* = 0.87). Follow-up comparisons revealed that stroking conditions (EMM = 4.37, SE = 0.42), collapsed across visual conditions, were perceived as more continuous than tapping conditions (EMM = −4.32, SE = 0.32; *z* = 13.92, *p* < .001). In addition, a significant interaction between tactile and visual conditions (*F*_1,30_= 239.31, *p* < .001, *d* = 0.36) revealed that the difference in continuity ratings between stroking and tapping increased from the photo conditions (V_P_T_Str_ - V_P_T_Tap_: *z* = 8.43, *p* < .001) to the conditions with touch videos (V_V_T_Str_ - V_V_T_Tap_: *z* = 16.96, *p* < .001, see Fig. 2A). The covariates age (*F*_1,28_ = 0.36, *p* = .55, *d* = 0.005) and gender (*F*_1,28_ = 0.43, *p* = .52, *d* = 0.02) did not yield significant effects.

**Fig. 2.**
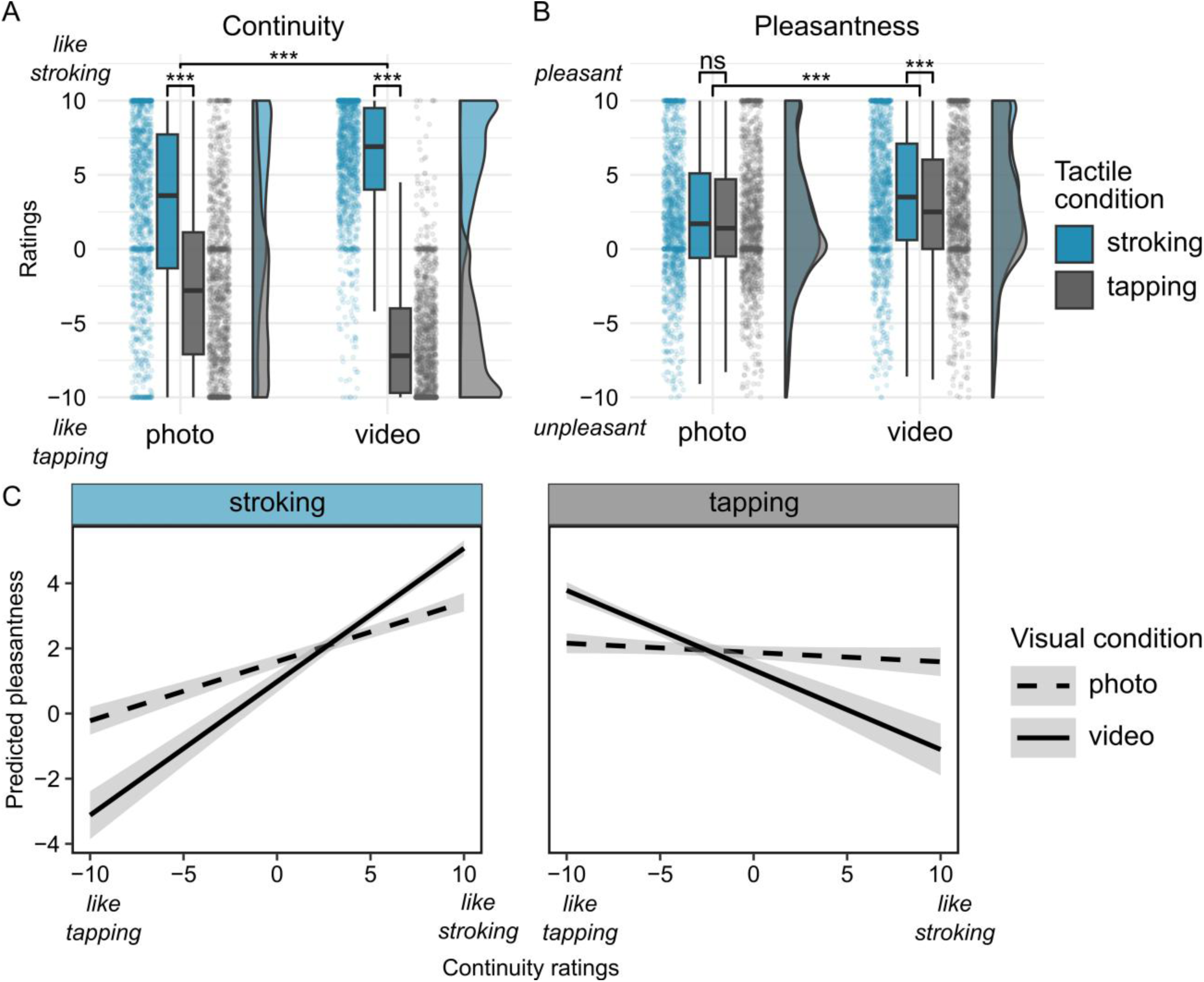
Continuity and pleasantness ratings showed differences between tactile stroking and tapping conditions as well as between video and photo conditions. Continuity ratings were higher in the stroking than in the tapping condition and this difference was larger when stroking and tapping were presented in the video condition (A). For pleasantness ratings, higher ratings were found in the video conditions than in the photo conditions. Furthermore, for tactile conditions, collapsed across visual conditions, stroking was perceived as more pleasant than tapping (B). Boxplots in A-B show the median (centre bar), first-to-third quartile (box), and the 1.5 inter-quartile range (whiskers). Single-trial data are shown as dots, and their distribution is illustrated as half-violin plots next to the boxplots. Post-hoc comparisons from linear mixed models are shown with brackets between individual or groups of boxplots. The significance of comparisons refers to marginal means as estimated from linear mixed models and not to the medians shown in the boxplots. (C) Regression slopes show the relationships between continuity ratings and predicted pleasantness for stroking (*left*) and tapping (*right*) in bisensory visual-photo (dashed line) and visual-video (solid line) conditions. The difference in slopes between video and photo conditions was significant for both stroking (difference = 0.17, SE = 0.05, *z* = 3.76, p< .001) and tapping (difference = −0.12, SE = 0.05, *z* = −2.68, p= .007), with the video condition amplifying the continuity-pleasantness relationship in both tactile conditions. Shaded areas represent the 95% confidence interval. ns: not significant (p > .05), *** p < .001

For pleasantness ratings, significant main effects of tactile conditions (*F*_1,30.44_ = 11.7, *p* = .002, *d* = 0.04) and visual conditions (*F*_1,30.0_ = 19.9, *p* = .001, *d* = 0.13) were observed. For the visual conditions collapsed across stroking and tapping, follow-up comparisons revealed higher pleasantness ratings in the video conditions (EMM = 3.08, SE = 0.61) than in the photo conditions (EMM = 1.93, SE = 0.61; *z* = 4.46, *p* < .001). Tactile conditions, collapsed across visual conditions, showed that stroking (EMM = 2.7, SE = 0.61) was perceived as more pleasant than tapping (EMM = 2.32, SE = 0.59; *z* = 3.42, *p <* .001). There was also a significant interaction between tactile and visual conditions (*F*_1,183.24_ = 7.23, *p* = .008, *d* = 0.03). To further explore the effects of visual conditions on stroking and tapping for pleasantness ratings, follow-up analyses of simple effects within each visual condition were conducted. In the touch video condition, stroking (EMM = 3.38, SE = 0.62) was rated as significantly more pleasant than tapping (EMM = 2.78, SE = 0.6; *z* = 4.13, *p* < .001). In contrast, no significant differences in pleasantness ratings were observed in the photo condition (stroking – tapping: *z* = 1.12, *p =* 0.26, see Fig. 2B). Additionally, as a control, the fixed effects for the covariates age (*F*_1,28.03_ = 2.03, *p* = .17, *d* = 0.06) and gender (*F*_1,28.03_ = 0.73, *p* = .4, *d* = 0.112) were both confirmed to be non-significant. In summary, stroking was perceived as more continuous and pleasant than tapping, especially when videos were presented alongside tactile stimulation.

Next, we examined the relationship between continuity and pleasantness ratings in an LMM. In this analysis, pleasantness ratings served as the outcome variable, continuity ratings, tactile and visual conditions as predictors, and gender and age as covariates. A three-way interaction showed that the relationship between continuity and pleasantness ratings varied across tactile and visual conditions (*F*_1,27.83_= 12.99, *p* = .001, *d* = 0.026, Fig. 2C). For stroking, higher continuity ratings predicted increased pleasantness. This relationship was stronger in the touch video condition (slope = 0.23, 95% CI [0.143, 0.321]) compared to the photo condition (slope = 0.063, 95% CI [0.015, 0.111]). Furthermore, tapping trials paired with touch videos were perceived as less continuous and more pleasant (slope = −0.108, 95% CI [-0.204, −0.011]). In contrast, tapping trials presented with photos showed no significant relationship between continuity and pleasantness (slope = 0.016, 95% CI [-0.044, 0.076]). Overall, continuity ratings were correlated with pleasantness ratings across both tactile conditions. This correlation was significantly strengthened by the addition of videos that provided information about the type of touch.

Regarding the embodiment measures, there was no difference in location ratings (i.e., the feeling that the real arm was perceived at the location of the virtual arm) between the touch types presented in the visual-video condition (overall mean = 4.08, SD = 1.35; *F*_1,28.0_ = 0.06, *p* = .81) nor between male and female participants (*F*_1,28.0_ = 0.005, *p* = .94) or age (*F*_1,28.0_= 0.08, *p* = .79). The ownership subscale, i.e., the feeling that the virtual arm was part of one’s body, also did not reveal any significant fixed effects (overall mean = 3.28, SD = 1.25; tactile conditions: *F*_1,28.0_= 0.17, *p* = .68; gender: *F*_1,28.0_= 0.16, *p* = .69, age: *F*_1,28.0_ = 0.21, *p* = .65). These embodiment ratings are consistent with those of previous experiments using visuotactile stimuli (e.g., Höfle et al., 2012, 2013). Furthermore, no differences in agency ratings (i.e., the feeling of controlling the virtual arm) were observed between touch types (overall mean = 2.09, SD = 1.16; *F*_1,28.0_= 0.04, *p* = .84) or between males and females (*F*_1,28.0_= 0.15, *p* = .7) or age (*F*_1,28.0_= 0.01, *p* = .92). On average participants correctly answered 84.5% (SD = 11.6%) of the attention questions asked after 6.25% of the trials. Taken together, these results validate that the experimental setup allowed participants to perceive their arm as being aligned with the virtual arm, and generally also made them feel as if they were looking at their real arm. Furthermore, these subjective ratings were neither influenced by touch type, gender or the age of participants.

### 3.2 Event-related potentials in sensorimotor cortex

As a validation, we first established that the digitally delivered tactile stimuli result in prominent ERP components over the contralateral sensorimotor cortex. Fig. 3 shows ERPs in response to bisensory visuotactile stimuli (stroking and tapping with a photo of an arm) at the ipsilateral sensorimotor cortex (electrode C3) and the contralateral sensorimotor cortex (electrode C4) for the visual-picture condition. The analysis focussed on the N2cc component from 0.18 to 0.28 s after tactile stimulation and revealed a significant effect of laterality (*F*_1,30.0_ = 7.83, *p* = .009, *d* = 0.3), which did not differ between stroking and tapping conditions (*F*_1,60.0_ = 0.2, *p* = .66, *d* = 0.02). Follow-up analysis revealed a more negative N2cc amplitude at the electrode contralateral (C4) (EMM = −1.13, SE = 0.24) compared to the electrode ipsilateral to the touch stimulation (C3) (EMM = −0.36, SE = 0.21; *t*_30_= −2.8, *p* = .009). Together these results indicated that the digitally delivered tactile stimuli elicited reliable sensorimotor ERPs.

**Fig. 3.**
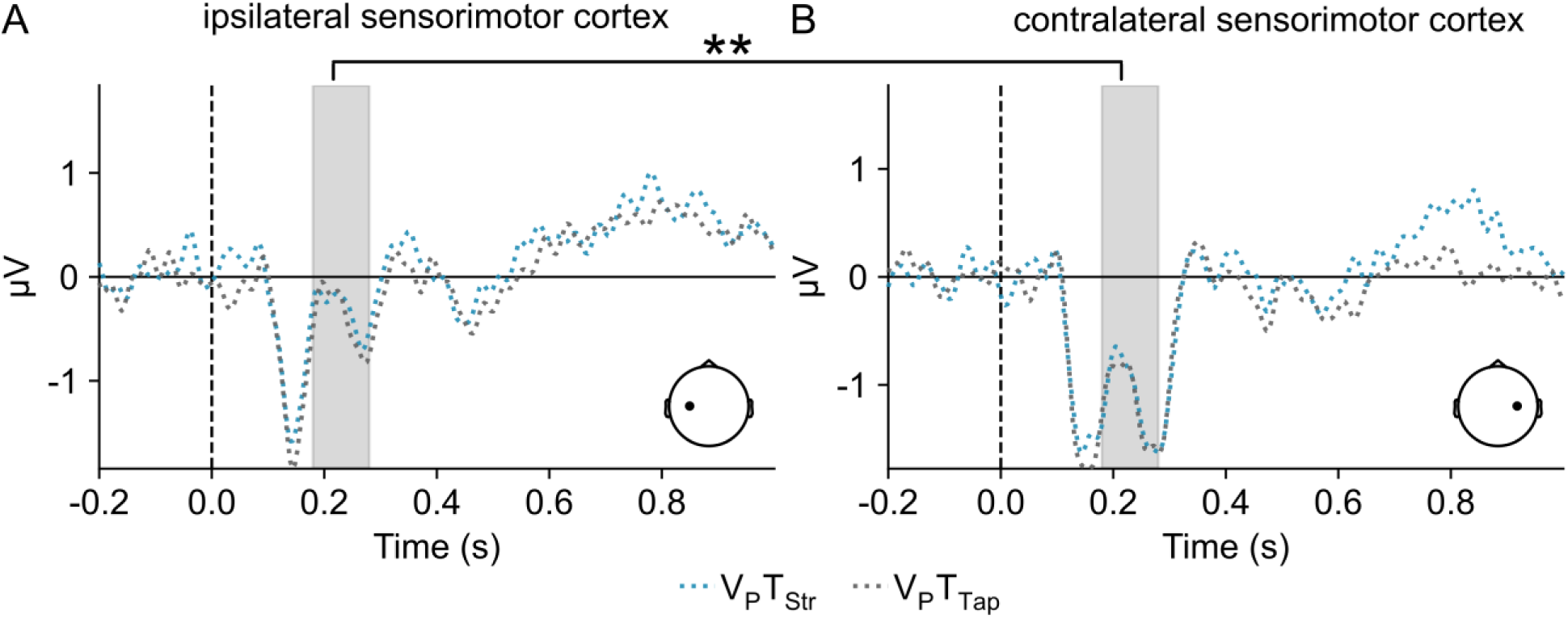
Event-related potentials to digitally delivered tactile stimuli over the ipsilateral and contralateral sensorimotor cortex. ERPs are plotted in the visual-photo condition for tactile stroking (V_P_T_Str_) and tactile tapping (V_P_T_Tap_) at ipsilateral sensorimotor cortex (electrode C3; panel A) and contralateral sensorimotor cortex (electrode C4; panel B). The negative ERP amplitudes of the N2cc component (0.18–0.28 s) were larger at the contralateral sensorimotor cortex than at the ipsilateral sensorimotor cortex (*t*_30_ = −2.8, *p* = .009). This demonstrates that digitally delivered tactile stimuli evoked sensorimotor ERPs. ** *p* < .01

### 3.3 Event-related potentials of cross-modal effects

Our main analyses focused on cross-modal effects of informative visual information on neural processing of tactile touch perception. Results of analysing ERPs from the stroking condition revealed one significant cluster from around 0.93 s to 1.34 s (p = .014) with a more prominent negative component for the video compared to the photo condition. This cluster encompassed seven left centroparietal sensors (Fig. 4A). In contrast, analysis of the tapping conditions did not reveal significant differences between video and photo conditions. The interaction contrast between the tactile and visual conditions revealed a significant cluster with negative t-values from about 1.63 to 1.88 s across six left centroparietal, two right central and two occipital electrodes (p = .037, Fig. 4B). Inspection of the time-series for the visual conditions revealed that the ERPs for stroking showed the opposite pattern (V_V_T_Str_ < V_P_T_Str_) compared to those observed for tapping (V_V_T_Tap_ > V_P_T_Tap_).

**Fig. 4.**
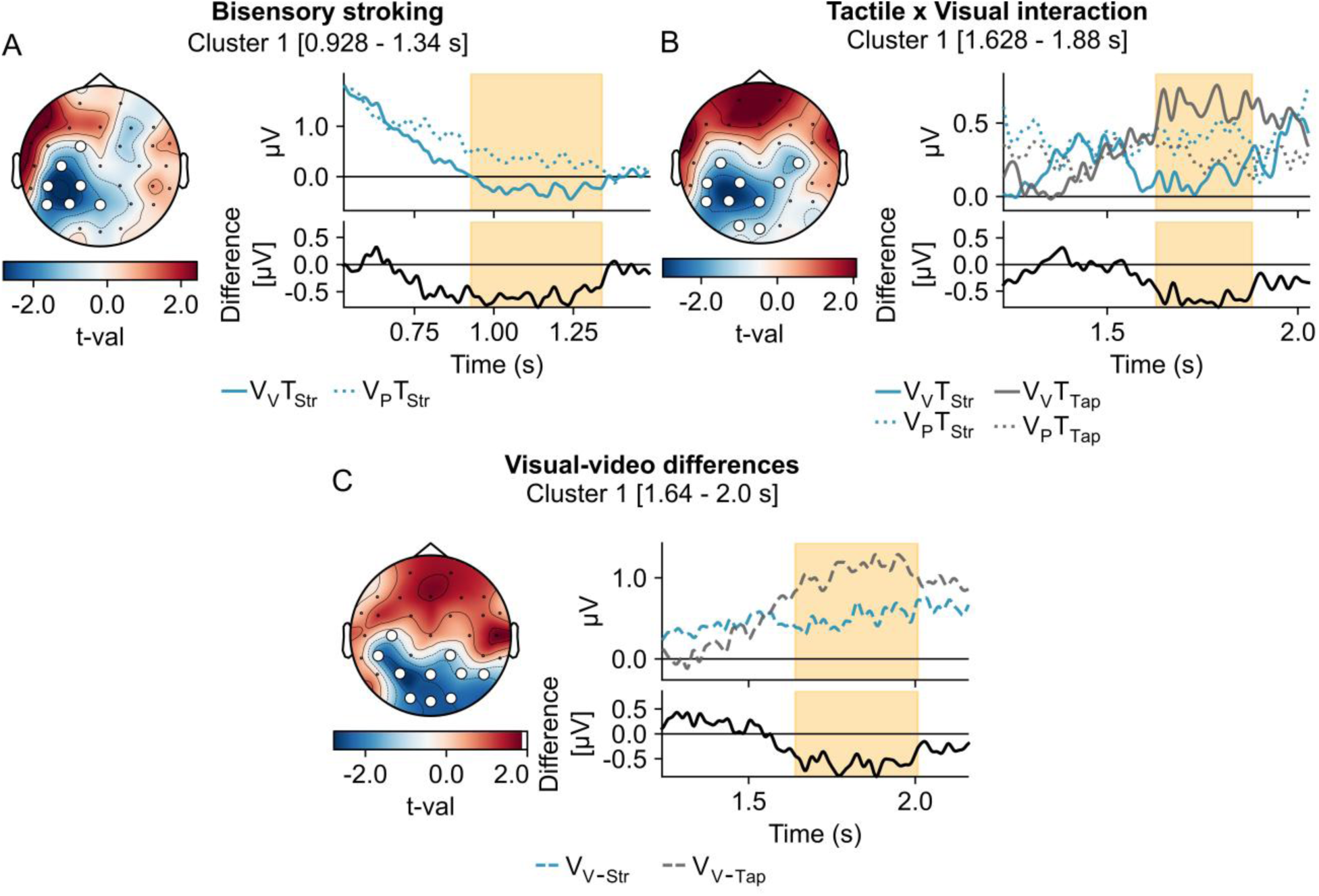
ERP differences between video and photo stimuli for the stroking condition, and for the interaction between tactile and visual conditions. (A) *Left*: A significant cluster comprising of seven left centroparietal electrodes was found between 0.93 and 1.34 s for the difference between the video (V_V_T_Str_) and photo stimuli (V_P_T_Str_) for the stroking condition. *Right*: Time course of the cluster amplitudes for the video and photo stimuli (top) and amplitude difference between the two conditions (V_V_T_Str_ - V_P_T_Str_, bottom). (B) *Left:* For the interaction between tactile and visual conditions, a significant cluster encompassing six left centroparietal, two right central and two occipital electrodes was observed between 1.62 and 1.88 s. *Right:* Time course of the cluster amplitudes for the four conditions (top) and amplitude difference for the interaction between tactile and visual conditions ((V_V_T_Str_ - V_P_T_Str_) – (V_V_T_Tap_ - V_P_T_Tap_), bottom). (C) *Left*: A significant cluster comprising of ten occipitoparietal electrodes was found between 1.64 and 2.0 s in the unisensory visual condition for the difference between stroking (V_V-Str_) and tapping (V_V-Tap_). *Right:* Time course of the cluster amplitudes of the visual-video conditions (top) and amplitude difference between both visual-video conditions ((V_V-Str_ - V_V-Tap_), bottom). These results show that cross-modal visual information affects neural processing of stroking and tapping at centroparietal clusters differently.

For an additional comparison, we examined differences between stroking and tapping in the unisensory video-only conditions (V_V_-_Str_ - V_V-Tap_) using a spatiotemporal cluster-based permutation test (two-tailed paired t-test, cluster-forming threshold = .05, 1000 permutations). A significant cluster emerged at around 1.64 s comprising ten occipitoparietal electrodes (p = .01, Fig. 4C). This indicates that processing differences in the unisensory visual condition coincided with the cluster found for the interaction contrast between tactile and visual conditions. Together, these results suggest that cross-modal visual information differently affects the processing of touch types (stroking vs. tapping) at centroparietal clusters.

### 3.4 Relations between cross-modal visual effects on ERPs and pleasantness ratings

Cluster-based permutation tests showed no significant correlation between ERPs and differences in pleasantness ratings between the two visual conditions (video versus photo) for stroking or for tapping. However, two significant right frontal clusters were observed for the positive correlation between the interaction contrasts of ERPs and pleasantness ratings (i.e., each computed as (V_V_T_Str_ - V_P_T_Str_) – (V_V_T_Tap_ - V_P_T_Tap_), Fig. 5A). One cluster was found between 1.376 and 1.432 s (p = .03), and another between 1.824 and 1.86 s (p = .042, Fig. 5B).

**Fig. 5.**
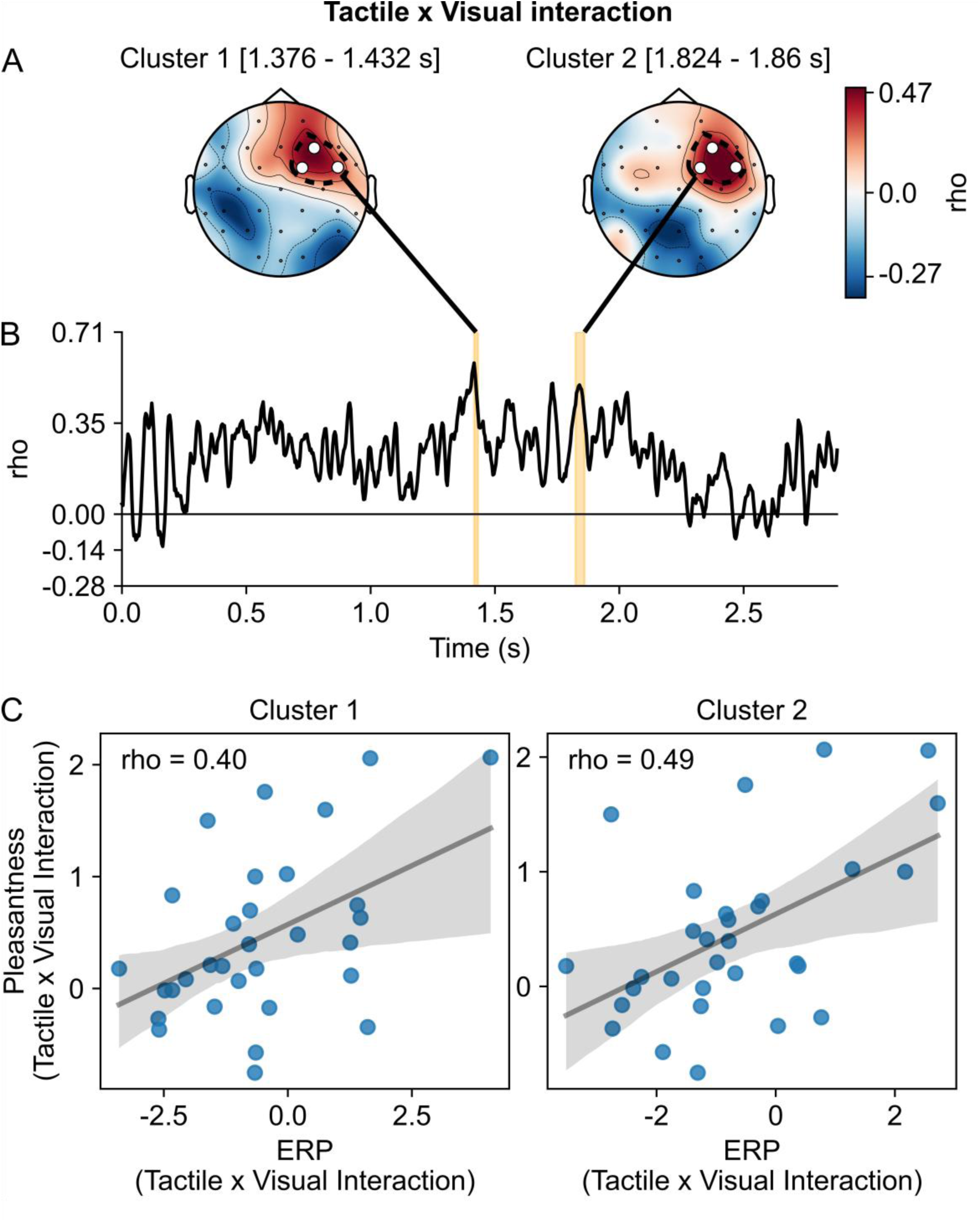
Clusters for the correlation between ERPs and pleasantness ratings as a function of the interaction between tactile and visual stimulation. (A) Two frontal right clusters were found for the interaction between tactile conditions (stroking vs. tapping) and visual stimulation (videos vs. photo). These clusters reflect positive correlations (Spearmans *rho*) between ERPs and pleasantness rating differences (i.e., (V_V_T_Str_ - V_P_T_Str_) – (V_V_T_Tap_ - V_P_T_Tap_)). In other words, participants who show greater cross-modal visual modulation (video > photo) for either stroking or tapping in frontal ERPs also demonstrate a corresponding cross-modal visual modulation of touch type in their pleasantness ratings. Electrodes of the significant clusters are highlighted as white dots. (B) Time course of average correlation across the three cluster electrodes is plotted with significant time-windows highlighted in orange. (C) Scatter plots serve the purpose to visualize the correlation between ERPs and pleasantness for the interaction between tactile and visual conditions at the participant level. Individual dots represent individual participants and their average difference ((V_V_T_Str_ - V_P_T_Str_) – (V_V_T_Tap_ - V_P_T_Tap_)) for ERPs and pleasantness ratings across significant sensors and time-windows for cluster 1 (*left*) and cluster 2 (*right*). Spearman’s rho is calculated and the best-fitting regression line is drawn with a 95% confidence interval to visualize the effects within both clusters.

These positive correlations indicate that an increase in the interaction between the tactile and visual conditions in ERPs is matched by an increase in pleasantness ratings for the same interaction term, i.e., (V_V_T_Str_ - V_P_T_Str_) – (V_V_T_Tap_ - V_P_T_Tap_). Therefore, the positive correlations demonstrate that participants who exhibit stronger cross-modal visual modulations (video > photo condition) for either stroking or tapping of frontal ERPs also exhibit the equivalent cross-modal visual modulation of touch types for pleasantness ratings (Fig. 5C).

## 4 Discussion

This study investigated how observing touch applied to a virtual arm that was aligned in orientation and location with the participants’ physical arm may affect the perception and tactile processing of digitally rendered stroking and tapping. Participants perceived stroking stimuli as more pleasant than neutral tapping stimuli. These differences were more pronounced in the video conditions, where visual information about the type of touch was provided. Late evoked brain responses reflecting cross-modal visual effects differed between the processing of the two touch types at centroparietal electrodes. Furthermore, cortical activity at frontal brain regions reflecting cross-modal effects on touch perception positively correlated with pleasantness ratings.

In terms of subjective ratings, differences in the perception of continuity were observed between stroking and tapping stimulations when touch was presented together with a photo of an arm. This result is also in line with those of a recent proof-of-concept study (Sousa et al., 2024). While for pleasantness some prior studies found stroking to be more pleasant than tapping (Etzi et al., 2018), and being associated with positive affective physiological responses (Etzi et al., 2018; Sousa et al., 2024), no differences in physiological markers of positive affect between stroking and tapping were also observed (McIntyre et al., 2022). In the present study, we also did not find differences in pleasantness ratings between touch types in the photo conditions. However, stroking was perceived as more continuous and more pleasant than tapping in the video conditions.

This suggest that the informativeness of cross-modal information may modulate pleasantness perception between touch types. Previous research has shown that tactile acuity improves when a photo of the stimulated body part is presented (Eads et al., 2015). Here, we extend this finding by comparing photos of an arm with videos containing information about touch type and movement for stroking and tapping. The spatiotemporally aligned videos of touch may make it easier to distinguish touch types and enhance the perception of pleasantness of the stroking stimulations. Supporting this, correlations between ratings of continuity and pleasantness were greater in the video than in the photo conditions for both touch types, suggesting that subjective perception of pleasantness could also be increased through cross-modal visual enhancement of tactile perception.

Analyses of the EEG data provided further insight into the cross-modal visual influences on touch processing. Comparing visual influence from videos versus photos for tactile perception of stroking showed differences at left centroparietal electrodes starting at around 0.92 s. This parietal involvement aligns with previous findings showing parietal cortex activation during visuotactile integration (Gentile et al., 2011; Pasalar et al., 2010). Additionally, multisensory integration in parietal areas was signalled by an increase in gamma power (Keil & Senkowski, 2018), which has been reported previously for congruent visuotactile stimuli (Kanayama & Ohira, 2009). Thus, the reported centroparietal cluster might facilitate cross-modal visual processing of touch movement information during pleasant touch. Another later centroparietal cluster resulted from the significant interaction effect between tactile conditions and cross-modal visual stimuli from around 1.62 s. This interaction resulted from differences in how cross-modal visual stimuli influence pleasant and neutral touch, and indicates a valence-dependent modulation of parietal, cross-modal related activity during pleasant touch.

This cluster for the interaction between tactile and visual conditions coincided with an occipitoparietal cluster exhibiting differences between unisensory visual conditions. Previously, unisensory visual differences were reported for processing of social versus non-social pictures (Schirmer & McGlone, 2019) and for videos of more pleasant versus less pleasant stroking (Morrison, Björnsdotter, et al., 2011). Furthermore, the occipitoparietal location of the cluster is partially aligned with parietal processing of visual socio-affective information (Masson & Isik, 2023). Therefore, we speculate that processing of unisensory visual valence, i.e., pleasant versus neutral touch videos, overlaps with and potentially contributes to cross-modal related activity differences between pleasant and neutral touch.

Further analyses examined the relationships between the cross-modal effects on ERPs and pleasantness ratings. These analyses revealed that the interaction effects observed for pleasantness ratings were correlated with the interaction effects observed for ERPs at two right frontal clusters, which had late onsets at 1.37 and 1.82 s post-stimulus.

Participants who exhibited larger ERP differences between visual information types for stroking versus tapping in frontal regions also exhibited corresponding patterns in pleasantness ratings. The spatiotemporal characteristics suggest a sustained late positive potential (Ackerley et al., 2013), which could reflect late affect-related valuation processing. These findings also align with those showing that reward-based value computations implicate the orbitofrontal cortex (Grabenhorst & Rolls, 2011). Since affective touch can be rewarding, these frontal activations may reflect reward processing (Rolls et al., 2003). Thus, our findings suggest that differences in frontal valuation induced by cross-modal visual information processing are associated with an enhanced perception of the pleasantness of pleasant touch relative to neutral touch.

In previous studies, the characteristic EEG features associated with the processing of pleasant touch have been identified for unisensory tactile stimulation. These features include the sN400 (Schirmer, Lai, et al., 2022; Schirmer & McGlone, 2019) and a late positive potential at around 0.7 s (Ackerley et al., 2013; Schirmer et al., 2023; Schirmer, Lai, et al., 2022; Schirmer & McGlone, 2019). The difference in latencies compared to our study may be attributed to differences in stimulus features. First, rather than directly comparing unisensory pleasant and neutral touch, we investigated how visual stimuli cross-modally affect different tactile conditions. Second, the relatively late (> 1.37 s) differences in cross-modal effects observed between the video conditions may be due to the similarity of the visual and tactile stimulation in the stroking and tapping conditions during the first 0.7 s (see Fig. 1C).

When interpreting the results, one has to consider that the digitally generated tactile stimuli were presented using a recently introduced arm-sleeve (Sousa et al., 2024). Although shape-memory alloy-based wearables have advantages in terms of flexibility and weight (Chernyshov et al., 2018), some limitations in the speed, duration and type of touch presentation (Boyraz et al., 2018) may restrict the use of this arm-sleeve in other experimental designs. For example, shorter pleasant and unpleasant visuotactile stimuli would allow for more direct comparisons of EEG patterns with studies on unisensory visual processing (Masson & Isik, 2023) or unisensory tactile processing (Schirmer, Lai, et al., 2022). Nevertheless, the results of our study, which were obtained by adding semantically meaningful visual information to affective touch, highlight potential avenues for future studies in presenting visuotactile stimuli with larger differences in valence and arousal (Spence, 2022).

In summary, our study shows that informative visual input enhances differences in pleasantness perception between continuous stroking and discrete tapping stimuli that were digitally generated through an arm-sleeve. This cross-modal enhancement effect is mediated by distinct neural processing recorded at centroparietal and frontal electrodes. While previous research has demonstrated that observing social touch activates areas involved in social perception processing, such as the posterior superior temporal sulcus (Masson & Isik, 2023), our study reveals that visually informative inputs cross-modally influence touch processing in centroparietal regions before valuating touch pleasantness. The valuation itself was associated with subsequent cross-modal visual modulation in frontal areas. Our findings establish that semantically congruent visual input influences perceived touch pleasantness. Considering the complexity of social interactions, future translational research could examine how wearable technology could compensate for the absence of physical contact in virtual reality (Fitzek et al., 2026; Li & Fitzek, 2023). For example, researchers could investigate how visually inputs from avatars cross-modally influence affective touch perception. This translational research would directly inform the development of new e-healthcare applications (McGlone et al., 2024).

## Declaration of competing interests

All authors declare no conflicts of interest.

## Data and code availability statement

All the processed data and analysis scripts are available upon request.

## Author contributions

**Brais Gonzalez Sousa:** Conceptualization, Data curation, Formal analysis (lead), Investigation, Methodology, Software, Visualization, Writing - original draft. **Daniel Senkowski:** Conceptualization, Methodology, Formal analysis (supporting), Writing - review & editing. **Shu-Chen Li:** Conceptualization, Funding acquisition, Methodology, Resources, Formal analysis (supporting), Writing - review & editing.

## Acknowledgments

We would like to thank Anna Weinert and Elitsa Boneva for their help with data collection. Brais Gonzalez Sousa and Shu-Chen Li acknowledge the financial support by the Federal Ministry of Research, Technology and Space of Germany in the programme of “Souverän. Digital. Vernetzt.”. Joint project 6G-life, project identification number: 16KISK001K. Shu-Chen Li is also funded by the German Research Foundation (DFG, Deutsche Forschungsgemeinschaft) as part of Germany’s Excellence Strategy – EXC 2050/1, 2050/2 – Project ID 390696704 – Cluster of Excellence “Centre for Tactile Internet with Human-in-the-Loop” (CeTI) of Technische Universität Dresden. Daniel Senkowski acknowledges support by the German Research Foundation (Project ID: SE1859/10-1).

## Notes

### Competing Interest Statement

The authors have declared no competing interest.

### Summary of Updates

Updated the title and abstract to better reflect the content of the paper; Updated some sections of the Introduction to better introduce into the topic of the paper; Moved sections from the Supplement to the Main document; Re-analysed EEG data due to applying average reference before channel interpolation; Updated results of behavioural and EEG data according to newly preprocessed EEG data; Added Fig.3 and Section 3.2 to validate sensorimotor ERPS; Updated Fig. 2-5 and Discussion to reflect updated results; Supplemental files updated.

