## Supplementary material for "Seeing touch enhances the perception and processing of digitized gentle stroking"

**Technical details on the shape-memory alloy (SMA) based arm-sleeve for touch actuation**

The SMA based arm-sleeve had thin wires knitted onto plasters (3 x 3 cm) that were arranged in a 3 (columns) by 5 (rows) structure on the arm-sleeve (Fig. 1B in main text). When the SMA arm-sleeve is activated, the wires contract row-by-row sequentially, thereby generating a gentle moving and squeeze-like sensation. The intensity of tactile stimulation was manipulated by the current passing through the wire, which in turn affects the degree of wire contraction and was set to be identical for both types of touch stimulations, i.e., 340 mA resulting in a 4% contraction equivalent to 1.44 N (for further details, see Muthukumarana et al., 2020).

The actuation of one row took 1 s and the digital stroking touch pattern was generated by consecutively actuating four rows of the SMA arm-sleeve (Fig. 1C in main text). Actuation-overlap defines the overlap between the actuations of successive rows so that the next row starts contracting 375 ms before the previous row finished actuating. This results in an actuated stroking velocity of roughly 4.17 cm/s, since four rows (covering a distance of 12 cm distance) are actuated in 2.875 s. Recently, we showed that this duration of actuation-overlap reliably generated a stroking sensation that elicited higher levels of positive affect compared to the tapping stimulation, as measured by facial muscle activity (Sousa et al., 2024). For the tapping stimulation, the first and fourth row were actuated with a delay of 0.875 s in between, which also results in the same trial duration of 2.875 s as for the stroking stimulation.

To reduce tactile habituation, the onset location and movement direction of the touch stimulations varied across trials. Specifically, four touch pattern variations were generated for each of the two touch types. Touch onset was randomized across two touch directions with onset at two distinct locations: wrist to elbow (stroking: row 1 to 4, or 2 to 5; tapping: row 1 then row 4, or row 2 then row 5), and elbow to wrist (stroking: row 5 to 2, or 4 to 1; tapping: row 5 then row 2, or row 4 then row 1).
